# The IMMUNE-ASSOCIATED NUCLEOTIDE-BINDING 9 protein is a regulator of basal immunity in *Arabidopsis thaliana*

**DOI:** 10.1101/277269

**Authors:** Yuanzheng Wang, Yansha Li, Tabata Rosas-Diaz, Carlos Caceres-Moreno, Rosa Lozano-Duran, Alberto P. Macho

## Abstract

A robust regulation of plant immune responses requires multitude of positive and negative regulators that act in concert. The immune-associated nucleotide-binding (IAN) gene family members are associated with immunity in different organisms, although no characterization of their function has been carried out to date in plants. In this work, we analyzed the expression patterns of *IAN* genes and found that *IAN9* is repressed upon pathogen infection or treatment with immune elicitors. *IAN9* encodes a plasma membrane-localized protein that genetically behaves as a negative regulator of immunity. A novel *ian9* mutant generated by CRISPR/Cas9 shows increased resistance to *Pseudomonas syringae*, while transgenic plants overexpressing *IAN9* show a slight increase in susceptibility. *In vivo* immunoprecipitation of IAN9-GFP followed by mass spectrometry analysis revealed that IAN9 associates with a previously uncharacterized C3HC4-type RING finger domain-containing protein that we named IAP1, for <u>I</u>AN9-<u>a</u>ssociated <u>p</u>rotein 1, which also acts as a negative regulator of basal immunity. Interestingly, neither *ian9* or *iap1* mutant plants show any obvious developmental phenotype, suggesting that they display enhanced inducible immunity rather than constitutive immune responses. Since both *IAN9* and *IAP1* have orthologs in important crop species, they could be suitable targets to generate plants more resistant to diseases caused by bacterial pathogens without yield penalty.

## Introduction

The plant immune system comprises an intricate network of receptors and regulators aimed at keeping the cellular homeostasis in the absence of pathogen threat and responding rapidly to biotic stimuli in order to prevent infection. Plants have evolved to perceive pathogen-derived molecules that constitute signals of a potential invasion, also called invasion patterns (Cook et al, 2015). Conserved microbial molecules constitute good targets for recognition by plants; some of these molecules have been shown to be perceived by plant cells as pathogen-associated molecular patterns (PAMPs; Boller and Felix, 2009). PAMPs are perceived at the cell surface by transmembrane pattern-recognition receptors (PRRs; Zipfel, 2014). PRRs act in coordination with several regulators and additional proteins that mediate signal transduction (Couto and Zipfel, 2016), including mitogen-activated protein kinases (MAPKs), calcium-dependent protein kinases (CDPKs), receptor-like cytoplasmic kinases (RLCKs), and respiratory burst oxidase homologs (RBOHs) (Macho and Zipfel, 2014; Bigeard et al, 2015). Downstream responses include the production of the phytohormone salicylic acid and antimicrobial compounds, the deposition of callose at the cell wall, and extensive transcriptional reprogramming (Boller and Felix, 2009). The activation of immunity is aimed at preventing the proliferation of the perceived pathogen, and prepares plant cells to mount an efficient defence response against subsequent biotic threats. However, defence is costly, in terms of energy investment and the concomitant disruption to the normal developmental program (Huot et al, 2014; Stael et al, 2015), and as such needs to be tightly regulated. For this reason, immune responses are inducible, and negative regulators ensure a firm control of their activated state. On the other hand, it has been demonstrated that activation of defence and inhibition of growth can be uncoupled, so that active defence and growth can occur simultaneously, indicating that developmental alterations are the consequence of an active process, and not necessarily of limiting resources (De Wit et al, 2013; Campos et al, 2016; Scheres and van der Putten, 2017).

Pathogens have developed strategies to manipulate plant cells in order to proliferate inside plant tissues. These include the suppression of plant immunity, the alteration of the physical environment, and the acquisition of nutrients to support their pathogenic lifestyle (Win et al, 2012). In Gram-negative bacterial pathogens, the most important virulence factor is the type-III secretion system (T3SS), which injects effector proteins directly into plant cells (type-III-secreted effectors, T3Es). The manipulation of plant cellular functions by T3Es is essential for bacteria to proliferate and is required for the development of disease (Macho and Zipfel, 2015; Macho, 2016). However, some plants have evolved intracellular receptors that can perceive T3E activities as an indication of pathogen invasion, hence becoming resistant to bacterial infection. These receptors contain nucleotide-binding and leucine-rich repeat domains (NLRs; Khan et al, 2016). NLR activation contributes to the development of defence responses similar to those established after PRR activation, but are often more intense, and sometimes lead to local cell death, which prevents further pathogen proliferation and spread (Chiang and Coaker, 2015). Both PRRs and NLRs constitute the basis of plant innate immunity, and are the major determinants of basal immunity in plants.

Guanosine triphosphate (GTP)-binding proteins are regulators of various biological processes in eukaryotic cells, such as signal transduction, cell proliferation, cytoskeletal organization, and intracellular membrane trafficking, and are classified into numerous families (Takai et al, 2001; Vernoud et al, 2003). IMMUNE-ASSOCIATED NUCLEOTIDE-BINDING/GTPases OF IMMUNITY-ASSOCIATED PROTEINS (IAN/GIMAP) proteins comprise a sub-family of GTPase-like proteins that has been found in anthozoans, vertebrates, and angiosperm plants (Nitta and Takahama, 2007; Weiss et al,2013). In vertebrates, proteins from the IAN/GIMAP family regulate the development and homeostasis of T cells and are associated with autoimmunity (Nitta and Takahama, 2007). The transcriptional regulation of genes encoding IAN/GIMAP proteins has been linked to immunity in different organisms: in mice, *IAN/GIMAP* genes are mostly expressed in immune tissues (Nitta et al, 2006), and they have also been reported as induced in corals after treatment with the bacterial immune elicitor muramyl dipeptide (MDP) (Weiss et al, 2013). In Arabidopsis, IAN/GIMAP proteins are defined by the presence of an avrRpt2-induced gene 1 (AIG1) domain, and contain conserved GTP-binding domains, including a P-loop motif known to bind GTP/GDP in Ras GTPases (Bourne et al, 1991), and coiled-coil motifs (Liu et al, 2008). Originally, *AtAIG1* (also known as *IAN8*) was defined as a gene overexpressed in response to the avirulent bacterial strain *Pseudomonas syringae* pv. *maculicola (Pma)* expressing the effector AvrRpt2 (Reuber and Ausubel, 1996). Additionally, computational analysis of transcriptomic data has revealed that the transcription of other *IAN* family members responds to different biotic stimuli: nematode infection induces the expression of *IAN3* and *IAN11*, while the transcription of *IAN11, IAN12* and *IAN13* is reduced by infection with *Myzus persicae* (Liu et al, 2008).

Despite the accumulating evidences that associate *IAN* genes with immunity in plants, no characterization of their function in this process has been carried out to date. Here, we analyzed the expression patterns of *IAN* genes and found that the expression of *IAN9* is repressed upon pathogen infection or treatment with immune elicitors. Further characterization indicated that *IAN9* encodes a plasma membrane-localized protein and that it genetically behaves as a negative regulator of immunity. *In vivo* immunoprecipitation of IAN9-GFP followed by mass spectrometry analysis revealed that IAN9 associates with a C3HC4-type RING finger domain-containing protein that we named IAP1, for <u>I</u>AN9-<u>a</u>ssociated <u>p</u>rotein 1. Interestingly, our results show that IAP1, like IAN9, negatively regulates immunity, raising the idea that these two proteins may work together in the control of immune responses.

## Results

### Expression analysis of the *IAN* gene family upon bacterial infection reveals differential expression patterns for *IAN7, IAN8*, and *IAN9*

The IAN protein family in Arabidopsis is composed of 13 members (Liu et al, 2008; Figure 1A). Phylogenetic analysis of IAN amino acid sequences shows a clear separation in different subgroups (Figure 1A, Figure S1). Previous reports have described changes in expression of *IAN* gene family members upon different biotic stimuli (Reuber and Ausubel, 1996; Liu et al, 2008). In order to characterize the transcriptional response of *IAN* genes to bacterial infection, we inoculated nine-to ten day-old Arabidopsis seedlings with the virulent pathogen *Pseudomonas syringae* pv. *tomato (Pto)* DC3000. Three *IAN* genes showed differential expression patterns upon bacterial infection: *IAN7* and *IAN8* showed a significant up-regulation, while *IAN9* showed a significant down-regulation (Figure 1B). We did not detect mRNA for *IAN1/2/4/6/10/11/12/13*, suggesting that these genes are not expressed in 10 day-old Arabidopsis seedlings in our experimental conditions. The pathogen-induced up-regulation of *IAN8* is reminiscent of the originally reported up-regulation of this gene by bacteria expressing the effector AvrRpt2, which induces activation of the plant NLR RPS2 (Reuber and Ausubel, 1996). Accordingly, we found that the differential expression of *IAN7/8* and *IAN9* also takes place upon infection of Arabidopsis rosette leaves with *Pto* expressing *AvrRpt2* (Figure 1C). The particular expression pattern of *IAN9* among the *IAN* gene family suggests an exclusive function for IAN9 rather than functional redundancy with other IAN family members. This idea is supported by the fact that *IAN9* constitutes a specific phylogenetic group within the *IAN* gene family (Figure 1A). For these reasons, we decided to focus our attention on IAN9.

**Figure 1.**
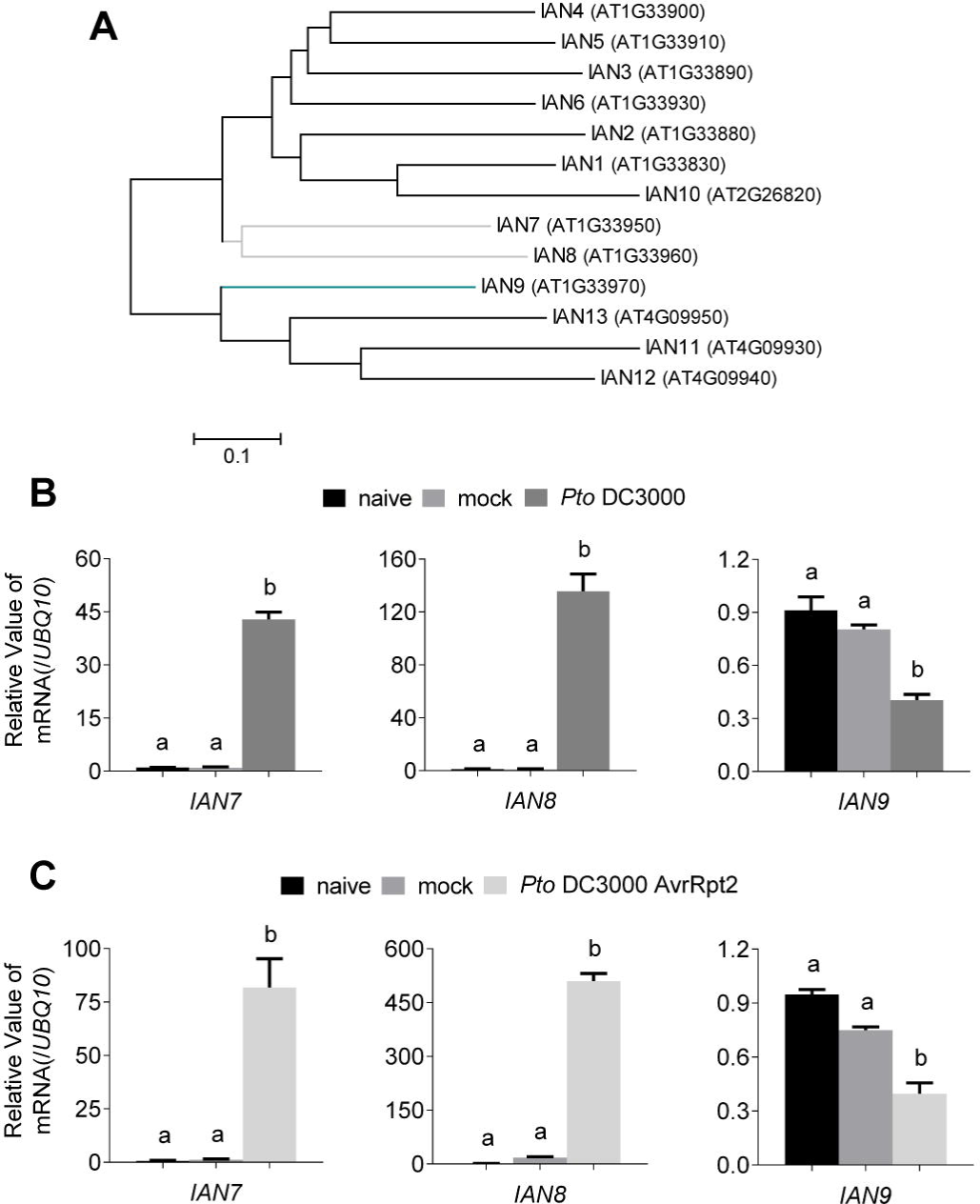
Transcription patterns of the *IAN* gene family in Arabidopsis.

(A) Phylogenetic tree of IAN proteins. The scale bar denotes a relative measure of evolutionary distance. (B and C) Relative expression of *IAN7, IAN8* and *IAN9* in 9-10 days-old Arabidopsis seedlings, 6 hours after inoculation with *Pto* DC3000 (B) or *Pto* DC3000 (AvrRpt2) (C). Real-time quantitative PCR results were normalized with *UBIQUITIN 10 (UBQ10*, AT4G05320). Values are means ± SEM (n=3 biological replicates; see methods). Statistical differences were calculated using one-way ANOVA (p<0.01). Each experiment was performed three times with similar results.

### *IAN9* expression is reduced upon chemical activation of plant immunity

*IAN9* is broadly expressed in cotyledons, hypocotyls, and roots of Arabidopsis seedlings (Figure S2). To dissect the bacteria-induced repression of *IAN9* transcription, we sought to determine whether perception of purified immune elicitors affects *IAN9* expression. For this purpose, we first treated Arabidopsis seedlings with the flagellin-derived peptide flg22, which is widely used as an elicitor of immune responses. Our results show that *IAN9* expression is significantly reduced one hour after flg22 treatment (Figure 2A). The perception of different invasion patterns, including flg22, leads to the production of the phytohormone salicylic acid (SA), which is a key player in the activation of plant immunity against biotrophic pathogens (Vlot et al., 2009). Treatment with SA for three hours led to a reduction of *IAN9* expression, although the abundance of *IAN9* transcripts returned to basal levels after a 6-hour SA treatment (Figure 2B), revealing a transient down-regulation of *IAN9* transcript abundance upon SA treatment. Finally, to determine whether the reduction on *IAN9* transcription upon flg22 treatment depends on the downstream SA accumulation (Tsuda et al., 2008), we performed flg22 treatment in the SA-depleted *sid2/NahG* plants, deficient in pathogen-induced SA biosynthesis (Wildermuth et al. 2001) and constitutively expressing the bacterial salicylate hydroxylase NahG, which degrades SA (Delaney et al., 1994). Interestingly, the flg22-triggered down-regulation of *IAN9* transcript abundance was also observed in *sid2/NahG* plants (Figure 2C), suggesting that SA is not required for the flg22-induced transcriptional repression of *IAN9.*

**Figure 2.**
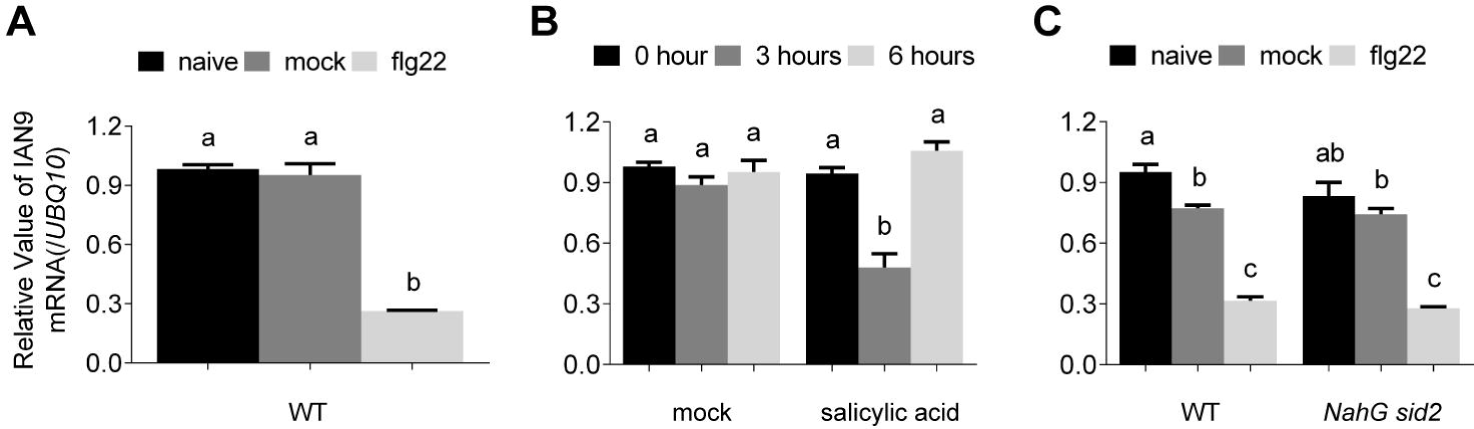
Treatment with flg22 or salicylic acid causes a reduction on the transcription of *IAN9.*

Relative expression of *IAN9* in 9-10 days-old Arabidopsis seedlings. Seedlings in (A) and (C) were treated with 100 nM flg22 for 1 hour. Seedlings in (B) were treated with 0.5 mM salicylic acid for 3 or 6 hours. Real-time quantitative PCR results were normalized with *UBIQUITIN 10 (UBQ10*, AT4G05320). Values are means ± SEM (n=3 biological replicates; see methods). Statistical differences were calculated using one-way ANOVA (p<0.01). Each experiment were performed three times with similar results.

### IAN9 localizes to the plasma membrane through its C-terminus

In order to investigate the subcellular localization of IAN9, we generated stable transgenic Arabidopsis lines expressing an N-terminal GFP-tagged IAN9 protein (see below for a detailed characterization of these lines), and used confocal microscopy to determine the localization of GFP-IAN9. Contrary to free GFP, which shows a nuclear/cytoplasmic localization, GFP-IAN9 specifically localized at the cell periphery (Figure 3A). This localization was similar to that observed for well-characterized plasma membrane (PM)-localized proteins, such as the brassinosteroid (BR) receptor BRI1 (Wang et al., 2001; Figure 3A). To determine whether GFP-IAN9 is associated to membranes, we used the lipophilic fluorescent dye FM4-64, which is rapidly incorporated into membranes upon contact with plant cells (Fischer-Parton et al., 2000; Bolte et al., 2004). Our results show that GFP-IAN9 fluorescence co-localizes with FM4-64-labeled compartments (Figure 3B and 3C), similar to BRI1-GFP fluorescence, and different from free GFP (Figure 3B and 3C). To further confirm that IAN9 localizes at the PM, we performed plasmolysis assays by treating plant tissues with 1 M NaCl. Upon plasmolysis, both GFP-IAN9 and BRI1-GFP were detected in Hechtian strands, which represent sites of incomplete PM retraction from the cell wall (Figure 3D). Altogether, our microscopy analysis indicates that IAN9 localizes at the PM in plant cells. The C-terminal domain is not conserved among IAN proteins (Figure S1). Interestingly, the C-terminal sequence of IAN9 shows an over-representation of positively charged amino acids (KKLRENLERAEKETKELQKKLGKCINL; 33.3% of R/K), not present in other IAN proteins (Figure S1 and S3A), which could mediate an interaction with the negatively charged phospholipids of the PM. To test this hypothesis, we generated Arabidopsis stable transgenic lines expressing a truncated GFP-IAN9 version lacking the 27 C-terminal amino acids (IAN9ΔC-27). The IAN9ΔC-27 version lost the specific PM localization seen for wild type GFP-IAN9, and was mostly detected in the cytoplasm (Figure S3B). Additionally, we found that, when GFP is fused to the C-terminal end of IAN9 (IAN9-GFP), this protein loses its specific PM localization, and is mostly found in the cytoplasm (Figure S3B). Compared to IAN9, the IAN8 C-terminus has a lower representation of positively charged amino acids (18.5%; Figure S3A), and we found that a N-terminal GFP-tagged IAN8 (GFP-IAN8) localizes to the cytoplasm (Figure S3C). Altogether, these results suggest that the exclusive C-terminal domain of IAN9 is required for its localization at the PM.

**Figure 3.**
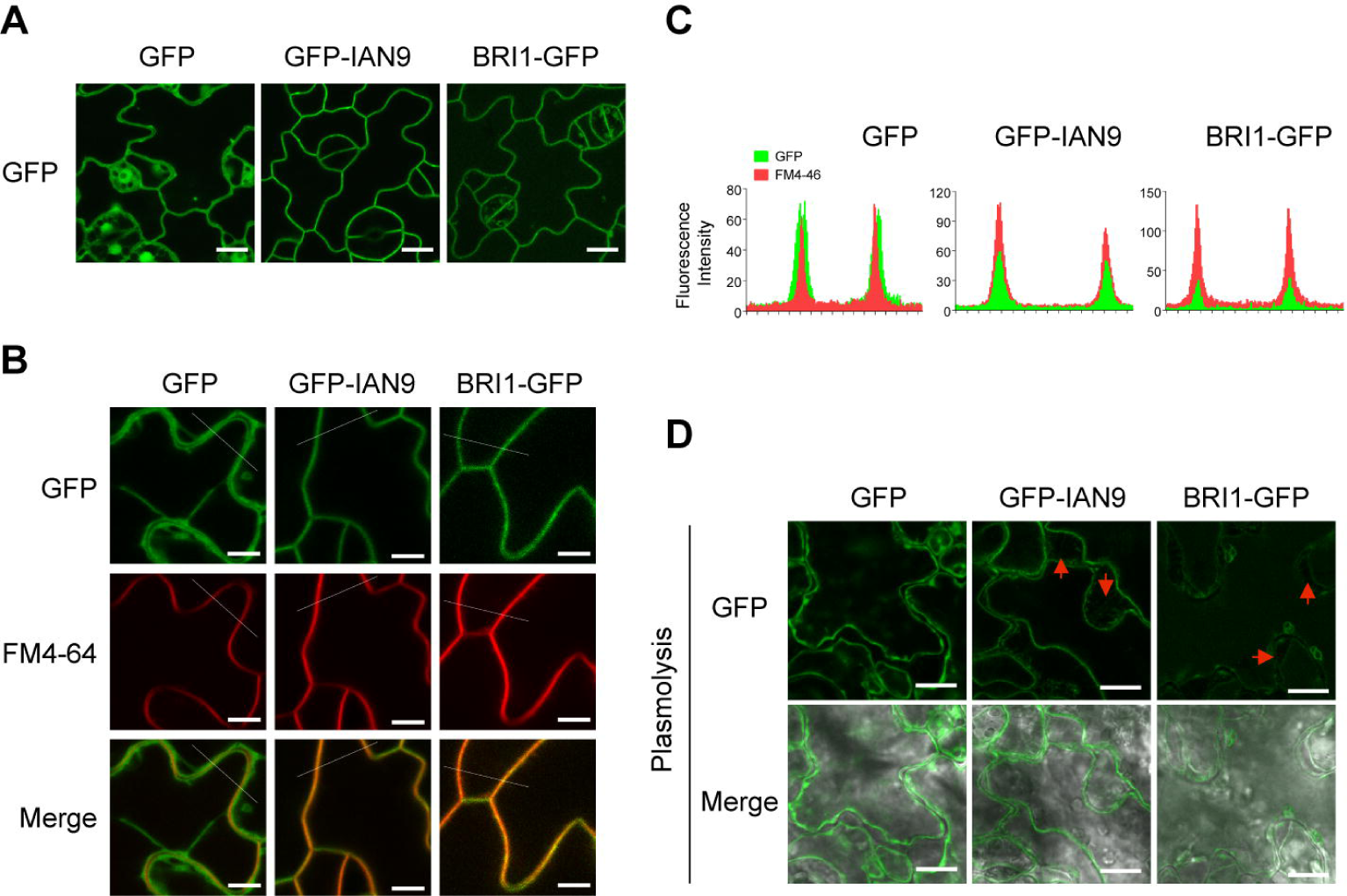
GFP-IAN9 localizes to the plasma membrane in *Arabidopsis*.

(A) Confocal images of GFP, GFP-IAN9, and BRI1-GFP in *Arabidopsis* cotyledon epidermal cells. Bar=10 μm. (B) Confocal images of GFP, GFP-IAN9, and BRI1-GFP in *Arabidopsis* cotyledon epidermal cells. *Arabidopsis* seedlings were treated FM4-64 for 5 minutes before confocal imaging, and the FM4-64 signal is shown in red. Bar=5 μm. (C) Fluorescence intensity through the thin lines shown in (B). (D) Confocal images of GFP, GFP-IAN9, and BRI1-GFP in *Arabidopsis* cotyledon epidermal cells after plasmolysis (5-minute treatment with 1M NaCl). Red arrows indicate the presence of Hectian strands. Bar=10 μm.

### Generation of *IAN9* knock-out and overexpression lines

Public repositories for Arabidopsis T-DNA insertion lines contain four independent lines with T-DNA insertions in the *IAN9* genomic locus: *SAIL_167_B02, SALK_534_B01, SALK_144369*, and *GK-146B08.* For all these lines, the insertion site is located in the predicted promoter region of *IAN9.* Among them, *GK-146B08* is the line harboring the nearest T-DNA insertion to the *IAN9* start codon (Figure S4A), and therefore we chose this line for further analyses. Sequencing analysis results showed that the insertion site is located 76 bp upstream of the 5’-UTR of the *IAN9* gene, and 319 bp upstream of the *IAN9* start codon (Figure S4A and S4B). RT-PCR and RT-qPCR analyses showed that this mutant has approximately a 3-fold reduction in the amount of *IAN9* transcripts (Figure S4C and S4D), indicating that this line is a knockdown *ian9* mutant. To further characterize this line, we determined the *IAN9* transcript levels upon bacterial inoculation. Surprisingly, we found that *IAN9* transcript levels increased significantly upon inoculation with *Pto* or *Pto* (AvrRpt2), reaching higher levels than those observed in wild type plants (Figure 1 and S4E). These findings indicate that the T-DNA insertion in the *IAN9* promoter generates an aberrant expression pattern of *IAN9* in this line, rendering it unusable for the functional characterization of this gene.

In order to perform genetic analysis of the contribution of IAN9 to plant immunity, we generated *ian9* mutant lines using CRISPR/Cas9-assisted genome editing (Feng et al, 2013; Mao et al, 2013). We selected the best predicted target site in the *IAN9* gene sequence for recognition by the Cas9/sgRNA complex (Figure S5), and performed the targeted mutagenesis as explained in the methods section. Sequencing of the resulting line showed an addition of 1 base pair in the second exon of *IAN9*, creating a premature stop codon 366 base pairs downstream of the start codon (Figure S6A), which generates a truncated protein with a disrupted GTP-binding domain. Upon selection of seedlings containing the *ian9* mutation in homozygosis, we selected a line in which the Cas9 gene was segregated out (Figure S6B); this Cas9-free *ian9* line was used for further experiments. Additionally, as mentioned before, we generated Arabidopsis transgenic lines overexpressing *IAN9* in a Col-0 wild type background, using a 35S promoter to express a *GFP-IAN9* fusion. We selected two independent homozygous lines that accumulated detectable amounts of GFP-IAN9 fusion protein (Figure S6C), and higher transcript levels of *IAN9* compared to those in Col-0 wild type (OE-*GFP-IAN9#3-10* and *OE-GFP-IAN9#7-1*) (Figure S6D). As controls, we selected two independent homozygous lines expressing free GFP, which did not show changes in *IAN9* transcription and accumulated detectable levels of free GFP (Figure S6C and S6D). Interestingly, although the 35S promoter led to high *IAN9* expression compared to Col-0 wild type, bacterial infection still caused a reduction in *IAN9* transcript levels (Figure S7), suggesting a post-transcriptional regulation of the abundance of *IAN9* transcripts. Overexpression of *IAN9* did not affect *IAN8* expression in basal conditions or upon bacterial infection (Figure S7).

### Plant growth and early immune responses are not affected by altered expression of *IAN9*

Both *IAN9* overexpressing or *ian9* knockout seedlings and adult plants were visually indistinguishable from the wild type (Figure S8 and S9). Given the predicted association of the IAN family to immune responses and the alteration in *IAN9* expression upon elicitation with flg22, we sought to determine whether IAN9 is involved in early PTI responses, namely the flg22-triggered burst of reactive oxygen species (ROS), and the activation of a cascade of mitogen-activated protein kinases (MAPKs) (Boller & Felix, 2009, Macho & Zipfel, 2014). Our results show that mutation or overexpression of *IAN9* did not have a detectable impact in the dynamics or total accumulation of ROS upon flg22 treatment (Figure S8B, S8C and Figure S10). Similarly, neither mutant nor overexpression (OE) lines showed differences in flg22-triggered MAPK activation compared to wild type plants or free GFP-expressing controls (Figure S8D and S8E). Taken together, these results suggest that alterations of *IAN9* expression do not affect early PTI responses.

### IAN9 negatively regulates plant immunity against *Pto* DC3000

To test whether IAN9 contributes to plant immunity against bacterial pathogens, we performed surface inoculation of the *ian9* mutant line with *Pto* DC3000 and determined bacterial replication in plant tissues. Our results show that knockout mutation of *IAN9* increased plant resistance against *Pto* DC3000, causing a 13-fold reduction on bacterial titers (Figure 4A). This increase in disease resistance does not seem to be caused by differences in basal expression of SA-dependent defense-related genes (Figure S11). In order to test whether the increase in resistance also occurs in a context of ETI, we syringe-infiltrated Arabidopsis rosette leaves with *Pto* DC3000 (AvrRpt2). However, no differences were detected in terms of replication of this strain (Figure 4B). To confirm that the observed increased resistance to *Pto* DC3000 is really due to the absence of *ian9*, we generated complementation lines expressing *GFP-IAN9*, driven by a 35S promoter, in the *ian9* knockout mutant background, where the GFP-IAN9 protein accumulated and localized to the PM (Figure S12A and S12B). Expression of GFP-IAN9 in the *ian9* background was able to rescue the level of growth of *Pto* DC3000 to that observed in wild type plants (Figure S12C), confirming the association of IAN9 with the observed increased resistance. Interestingly, transgenic lines overexpressing GFP-IAN9 in a wild type background showed a tendency to support higher bacterial loads compared to wild type or GFP-expressing lines, suggesting that IAN9 overexpression suppresses plant immunity against *Pto* DC3000, although such tendency was not always reproducible or statistically significant across eight independent biological repeats (Figure S13). This trend was not observed when plants were inoculated with a hypo-virulent DC3000 derivative unable to produce the virulence factor coronatine *(Pto* DC3000 COR-; Figure S14) (Melotto et al, 2006).

**Figure 4.**
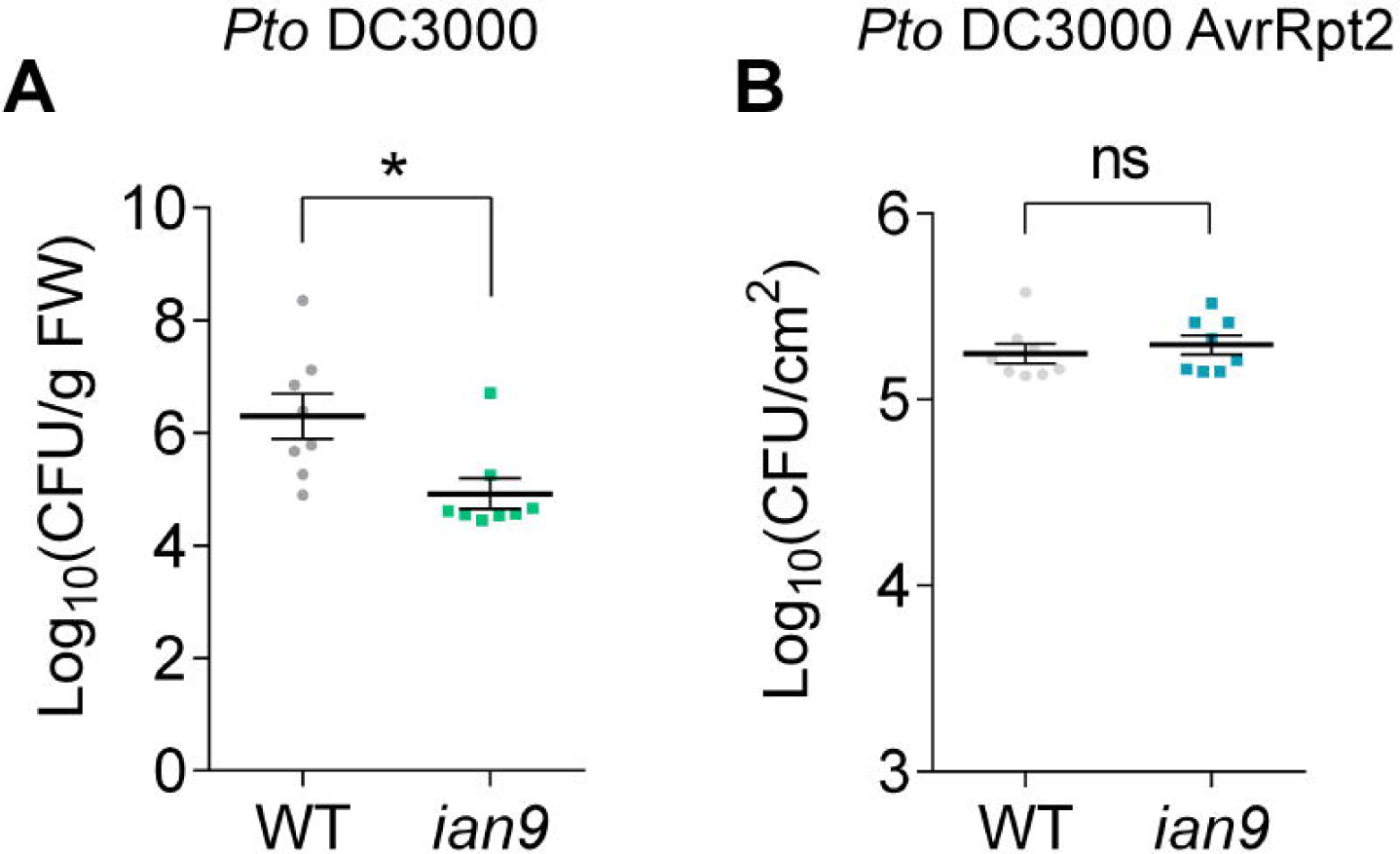
IAN9 negatively regulates plant immunity against *Pto* DC3000.

(A) Growth of surface (spray)-inoculated *Pto* DC3000 (0D600=0.1) in wild-type (WT) Col-0 and *ian9* mutant plants, 3 days post-inoculation (dpi). Experiments repeated more than three times with similar results. (B) Growth of *Pto* DC3000 (AvrRpt2) (0D_600_=0.001) infiltrated with a needleless syringe into wild-type (WT) Col-0 and *ian9* mutant plants, 3 days post-inoculation (dpi). Experiments performed twice with similar results. (A and B) Data were represented as means ± SE (n=8 independent plants). Statistical differences were calculated using a Student’s t-test. “ns” indicates no significant difference, and asterisk indicated significant difference (p<0.05).

### Identification of IAN9-interacting proteins

To characterize further the mode of action of IAN9, we searched for proteins physically associated with IAN9 in plant cells using Arabidopsis seedlings expressing *GFP-IAN9* (line *OE-GFP-IAN9#3-10*) and seedlings expressing free GFP as control. Upon GFP immunoprecipitation (IP) using agarose beads coupled to an anti-GFP nanobody (GFP-Trap beads), we detected associated proteins using liquid chromatography coupled to tandem mass-spectrometry (LC-MS/MS). In order to detect potential dynamic interactions, we additionally treated seedlings for one hour with flg22 or SA before immunoprecipitation. Two independent biological replicates showed a large number of proteins physically associated with IAN9, which we filtered using the following criteria: 1. Presence in both biological replicates; 2. Detection of two or more exclusive unique peptides; 3. Absence in the GFP control. After applying these filters, a total of 14 proteins were identified as IAN9 candidate interactors (Table S1).

Among these candidate interactors, we drew our attention to an uncharacterized protein encoded by the AT1G18660 gene (Table S1), which we named IAP1 for IAN9-ASSOCIATED PROTEIN1. Although our experimental approach does not provide a quantitative assessment of protein interactions, we noticed that the number of detected IAP1 peptides decreased in samples treated with SA, while it did not change substantially in samples treated with flg22 (Table S1). Domain analysis predicts the presence of three tetratricopeptide-like helical (TPR) domains, a C3HC4-type RING finger domain, and an ATP-dependent protease La (LON) domain in this protein (Figure S15A). Upon Agrobacterium-mediated transient expression in *Nicotiana benthamiana*, a GFP-IAP1 fusion protein localized at the cell periphery, nucleus (weakly), and around the nucleus; the latter most likely corresponds to the usual endoplasmic reticulum localization of PM-localized proteins overexpressed in this system (Figure S15B and S15C). Coexpression with RFP-IAN9 showed that both proteins co-localize upon transient expression (Figure 5A). To confirm the interaction between IAN9 and IAP1, we performed a targeted co-IP using GFP-IAP1 and an N-terminal fusion of IAN9 to the C-terminal domain of luciferase (Cluc-IAN9). Co-IP assays show that GFP-IAP1 interacts with Cluc-IAN9, but not with Cluc alone (Figure 5B); Cluc-IAN9 does not interact with free GFP (Figure S16). To determine whether the IAN9-IAP1 interaction is direct, we performed a luciferase complementation assay, transiently co-expressing Cluc-IAN9 and IAP1 fused to the N-terminal domain of luciferase (IAP1-Nluc). A positive control co-expressing AtSGT1b-Nluc and Cluc-AtRAR1 showed a strong luciferase signal (Figure S17A), as described before (Chen et al., 2008). On the contrary, tissues co-expressing Cluc-IAN9 and Nluc-IAP1 did not show any detectable luciferase signal (Figure S17A), although both proteins accumulated (Figure S17B). As an alternative technique to detect direct interactions, we employed FRET-FLIM by co-expressing GFP-IAP1 and RFP-IAN9. However, no difference in GFP fluorescence lifetime was detected when compared to control samples (Figure S17C). Although an influence of the tags cannot be ruled out, these results suggest that the interaction observed for IAP1 and IAN9 is most likely indirect.

**Figure 5.**
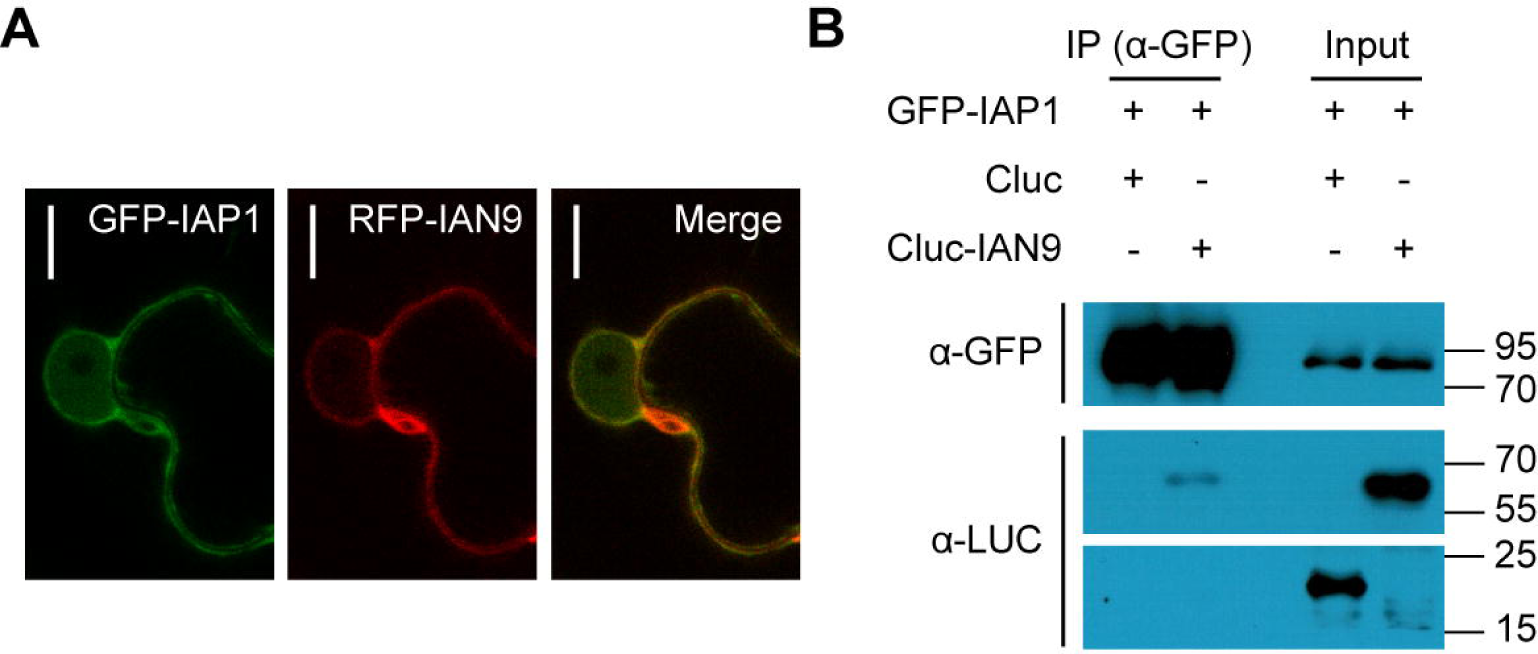
IAP1 interacts with IAN9 in *N. benthamiana* leaves.

(A) Confocal microscopy images showing the co-localization of GFP-IAP1 and RFP-IAN9 in *N. benthamiana* leaves. Bar=10 μm. (B) Cluc or Cluc-IAN9 was co-expressed with GFP-IAP1 in *N. benthamiana* before immunoprecipitation using GFP-trap beads. Immunoblots were analysed using anti-luc or anti-GFP antibody. Molecular weight (kDa) marker bands are indicated for reference. The experiments were repeated three with similar results.

### IAP1 negatively regulates plant immunity against *Pto* DC3000

Public repositories for Arabidopsis T-DNA insertion lines contain two independent lines with T-DNA insertions in the *IAP1* genomic locus: *SALK_119114* and *SALK_093498* (Figure S18A). We isolated homozygous lines containing these insertions (Figure S18B) and confirmed the absence of *IAP1* transcripts (Figure S18C), naming these lines *iap1-1* and *iap1-2*, respectively (Figure S18A). Both lines displayed wild-type-like growth and development when grown on soil under short-day conditions (Figure 6A), although mutant seedlings showed a slightly reduced root length compared to Col-0 wild type when grown in vertical MS plates (Figure S19).

To determine whether mutations in *IAP1* have an impact on plant resistance against bacterial pathogens, we performed surface inoculation of the *iap1* mutant lines with *Pto* DC3000. Our results show that both *iap1-1* and *iap1-2* mutant lines are more resistant than Col-0 wild type against *Pto* DC3000, showing a 19-fold reduction on bacterial loads (Figure 6B). However, none of these lines showed differences after inoculation with *Pto* expressing AvrRpt2 (Figure 6C). This enhancement of disease resistance did not seem to be caused by differences in basal expression of SA-dependent defense-related genes (Figure S11). Interestingly, these results resemble those obtained during the characterization of the *ian9* knockout mutant line (Figure 4), suggesting that both proteins are involved in the negative regulation of basal immunity against bacterial pathogens, and may act in the same pathway through physical association.

**Figure 6.**
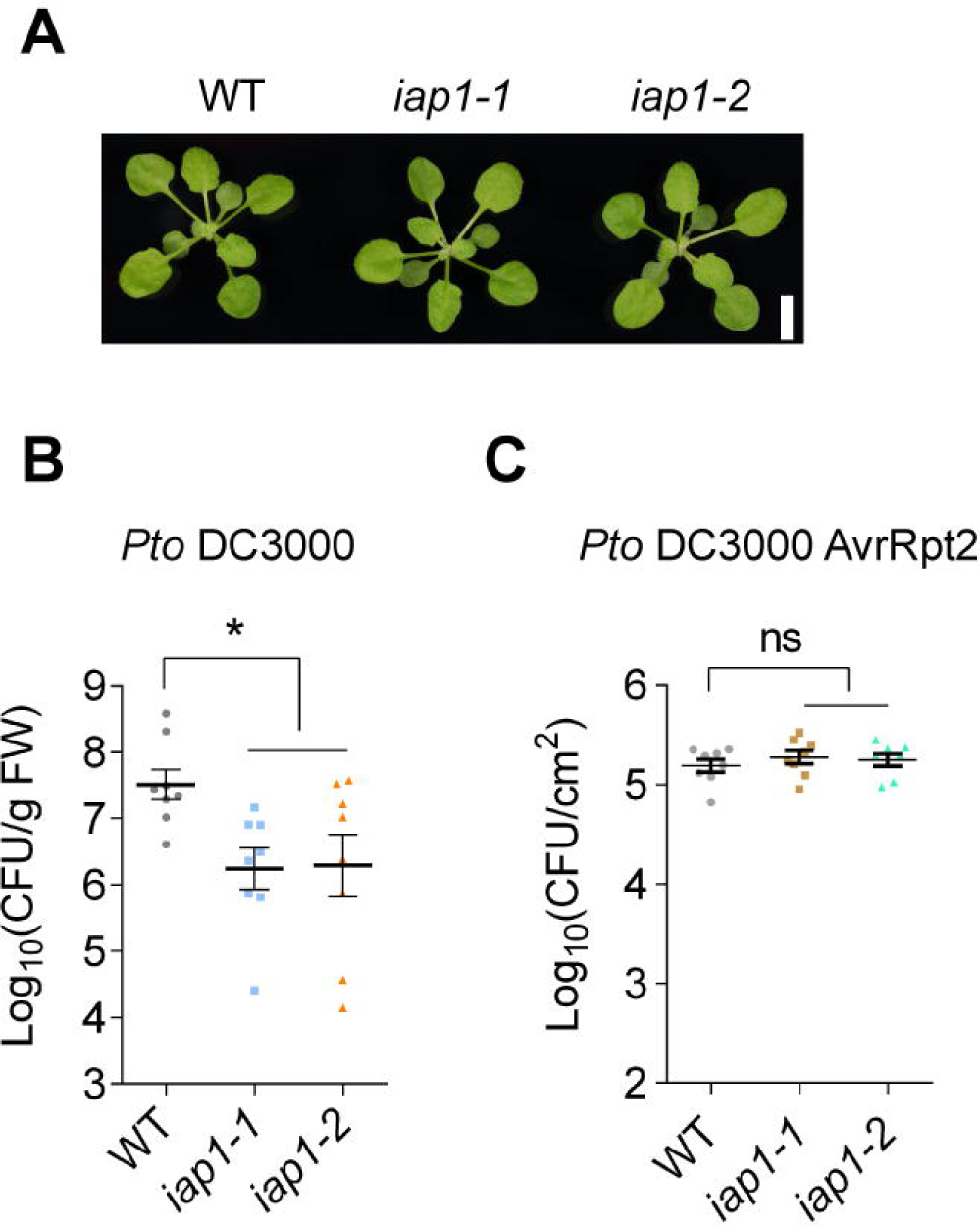
IAP1 negatively regulates plant immunity against *Pto* DC3000.

(A) Photography of four week-old Col-0 wild type (WT), *iap1-1*, and *iap1-2* plants, grown at a 8 h light/16 h dark photoperiod. Scale bar is 0.5 cm. (B) Growth of surface (spray)-inoculated *Pto* DC3000 (0D_600_= 0.1) in wild-type (WT) Col-0, *iap1-1*, and *iap1-2* mutant plants, 3 days post-inoculation (dpi). Experiments repeated more than three times with similar results. (C) Growth of *Pto* DC3000 (AvrRpt2) (0D_600_=0.001) infiltrated with a needleless syringe into wild-type (WT) Col-0, *iap1-1*, and *iap1-2* mutant plants, 3 days postinoculation (dpi). Experiments performed twice with similar results. (B and C) Data were represented as means ± SE (n=8 independent plants). Statistical differences were calculated using a Student’s t-test. “ns” indicates no significant difference, and asterisk indicated significant difference (p<0.05).

## Discussion

In this work, we characterized the transcription patterns of *IAN* family members in Arabidopsis upon infection with the bacterial pathogen *Pto* DC3000. *IAN7* and *IAN8*, which seem to form a separate phylogenetic subgroup, are up-regulated upon bacterial infection. On the contrary, *IAN9* seems to form a specific subgroup and shows a unique down-regulation upon bacterial infection. Infection with a virulent bacterial pathogen, such as *Pto DC3000*, triggers changes in plant cells caused by bacterial virulence activities (e.g. alteration of plant processes by T3Es) and basal immune responses. Therefore, formally, transcriptional changes upon infection could be associated with bacterial virulence or plant immunity. In this case, the down-regulation of *IAN9* expression upon bacterial infection seems associated with the activation of immunity, since we detected similar expression patterns upon treatment with elicitors of immunity, namely flg22 and SA. Interestingly, we detected a reduction in the amount of *IAN9* transcripts after bacterial infection even in transgenic lines where *IAN9* overexpression is driven by a constitutive 35S promoter, suggesting a post-transcriptional regulation of *IAN9* transcript abundance. Although the perception of flg22 leads to the production of SA (Tsuda et al., 2008), our data show that the flg22-triggered reduction of *IAN9* transcription takes place in *sid2/NahG* plants, indicating that this transcriptional change is independent of the observed SA-triggered reduction of *IAN9* transcription. SA-deficient mutants are partially affected in flg22-triggered induction of defence-related genes, probably caused by lower levels of the flg22 receptor FLS2 (Yi et al.,2013). Our results indicate that these lower levels of FLS2 in *sid2/NahG* plants are nevertheless sufficient to trigger the *IAN9* transcriptional repression, suggesting that *IAN9* transcription may be highly sensitive to external biotic stimuli.

Besides showing different expression patterns, the subcellular localization of IAN9 (mostly at the PM) is also different from that of IAN8 (mostly in the cytoplasm). This differential localization further suggests distinct functions for these two IAN family members. The C-terminal domain of IAN9, required for PM localization, is not present in other IAN proteins. This could indicate that the IAN9 mechanism for PM attachment is exclusive within the IAN family, although other IAN proteins could localize at the PM through different means.

Mutation of *IAN9* causes increased resistance to *Pto* DC3000, and this phenotype is rescued in complementation lines. It is worth considering the possibility that IAN9 is guarded by NLRs as it has been shown for some regulators of immunity (Khan et al, 2016), therefore explaining the increased resistance in the *ian9* knockout mutant. However, the absence of developmental phenotypes or constitutive immune responses makes this possibility unlikely. Additionally, overexpression lines show a tendency for increased susceptibility, although this phenotype was not always significant or reproducible, and this trend was not observed upon inoculation with a hypovirulent *Pto* DC3000 derivative strain. The lack of robustness in this phenotype could suggest that *IAN9* transcript abundance is not rate limiting; however, it could also be due to the potential post-transcriptional regulation of *IAN9* transcript abundance that we have seen in the *IAN9* overexpression lines, which show a reduction in the amount of *IAN9* transcripts after bacterial infection. The transient nature of *IAN9* transcriptional repression upon SA treatment, returning to basal levels after 6 hours, may indicate that the negative regulation exerted by IAN9 could be required to contribute to the repression of immune responses after immunity has been established.

Using IP followed by LC-MS/MS, we found IAP1 as a protein associated with IAN9. IAP1 contains three tetratricopeptide-like helical (TPR) domains, a C3HC4-type RING finger domain, and an ATP-dependent protease La (LON) domain. Upon transient expression, IAP1 co-localizes with IAN9 in *N. benthamiana* cells. Genetic analysis indicates that both IAN9 and IAP1 negatively regulate immunity against *Pto* DC3000. Given their physical association, it is possible that both proteins belong to a protein complex involved in the negative regulation of basal immunity against bacterial pathogens. Although we confirmed the association between IAN9 and IAP1 using targeted IP, we failed to detect a direct physical interaction using luciferase complementation or FRET-FLIM assays. Specific limitations of interaction techniques may be hindering the detection of a direct interaction; however, these data may also indicate that the interaction between IAN9 and IAP1 is indirect, perhaps mediated by other scaffolding components in the same protein complex. The Arabidopsis Interactome Database (http://interactome.dfci.harvard.edu/Athaliana/index.php) has reported several interactors for IAP1 in Y2H assays (Table S2). Interestingly, several of these interacting proteins are transcription factors, and others have predicted nuclear localization or undergo nucleo-cytoplasmic re-localization associated to their activity (Table S2). Considering that IAP1 interacts with several proteins with nuclear activities, it is tempting to speculate that the IAN9/IAP1 complex could associate with transcriptional regulators in the absence of biotic stress, acting as negative regulators of immune responses, and the complex may dissociate upon pathogen infection to allow for the activation of executor immune responses that restrict pathogen proliferation. This hypothetical model is in agreement with the observation that IAN9 does not seem to regulate early elicitor-induced responses (namely flg22-triggered ROS burst and MAPK activation). Given that *ian9* or *iap1* mutants do not show auto-immunity phenotypes, the final activation of immune responses may require additional activation by other immune regulators, of which the activity could be facilitated by the absence of IAN9 or IAP1.

The identification of sustainable sources of resistance against plant pathogens is essential to minimize crop losses due to diseases. A well-established approach consists on the mutation of negative regulators of immunity, or the so-called susceptibility genes, although this often leads to fitness costs or auto-immunity phenotypes, which obstruct their applicability in agriculture. Disease resistance based on loss-of-function mutations in *Mildew resistance locus o* (*Mlo*) genes is one of the best-known examples of this approach (reviewed in Kusch & Panstruga, 2017). However, *mlo* mutations sometimes have pleiotropic effects that may affect plant yield and increase susceptibility to other pathogens (Kusch & Panstruga, 2017). Altogether, our results reveal *IAN9* and *IAP1* as suitable targets for biotechnological approaches to generate crops with increased disease resistance to bacterial pathogens, since both *IAN9* and *IAP1* have orthologs in agriculturally important crop species (Figure S20 and S21). Interestingly, IAN9 and IAP1 behave as negative regulators of basal immunity, since *ian9* and *iap1* knockout plants show increased resistance to bacterial infection, but they do not show obvious differences compared to wild-type plants in terms of size or development. It remains to be determined whether the mutation of *IAN9* or *IAP1* orthologs in other plant species will have an impact on the plant response to other biotic or abiotic stresses. Alternative strategies to engineer resistance to plant diseases consider the pathogen-induced transcriptional and translational control of immune regulators (Gurr & Rushton, 2005; Xu et al, 2017). These strategies show that it is possible to generate plants where the expression of defence-related genes is restricted to cells undergoing pathogen attack, thus avoiding side effects on plant fitness. Current regulations hinder the use of transgenic plants to generate disease-resistant crops. Our *IAN9* mutagenesis approach shows that it is feasible to design CRISPR/Cas9-mediated strategies to obtain stable non-transgenic mutant plants with increased resistance to pathogens and no obvious developmental defects, paving the way to a potential application in breeding for disease resistance.

## Materials and Methods

### Plant materials and growth conditions

Arabidopsis seeds were sterilized with bleach solution (20% bleach and 0.1% Triton X-100) for 2-3 minutes, then washed with sterile water 4-5 times and sown on solid ½ MS medium (2.21 g Murashige & Skoog Basal Medium with Vitamins, 15 g Sucrose, 7 g Agar. 1 L). After stratification at 4°C for 3 days, the plates were transferred to a growth chamber (22°C, 16 h light/8 h dark) for germination and growth. For experiments involving mature Arabidopsis plants, sterile seeds were firstly stratified at 4°C for 3 days, then transferred to soil. Plants for bacterial infection assays were cultivated in a short day chamber (22°C, 8 h light/16 h dark photoperiod, 65% humidity). Plants for *Agrobacterium* transformation were grown in a long day growth room (22°C, 16 h light/8 h dark photoperiod).

### Bacterial infections

*Pseudomonas syringae pv tomato* DC3000 *(Pto* DC3000) and DC3000 (AvrRpt2) were streaked on selective ½ salt LB plates (10 g Tryptone, 5 g Yeast Extract, 5 g NaCl, 15 g Agar per 1 L) and cultivated at 22°C for 2 days. For *Pto* DC3000, bacteria were collected from the plates into sterile water, the OD was adjusted to a value of 0.1 (5×10^7^ cfu/ml) or 0.2 (10^8^ cfu/ml), and silwet-L77 was added to a final concentration of 0.03% before performing spray inoculation onto 3-4 week-old soil-grown Arabidopsis under short day conditions. The plants were then covered with cling wrap for 3 hours. The whole plants were harvested in 1.5 ml microcentrifuge tubes and weighed. For *Pto* DC3000 (AvrRpt2), the OD was adjusted to 0.1 (5×10^7^ cfu/ml) or 0.001 (10^5^ cfu/ml). The bacterial suspension was pressure-infiltrated into 5-6 week-old short day-grown Arabidopsis leaves with a needleless syringe (3 leaves per plant), and leaf discs were collected into 1.5 ml tubes using a leaf punch at three days post-inoculation. In both cases, plant tissues were ground and homogenized in sterile water before plating serial dilutions on selective Y salt LB agar plates. The plates were placed at 28°C for 1.5 days and the bacterial growth was calculated as colony-forming units.

For gene expression assays, sterile seeds were sown on ½ solid MS medium to germinate and grown for 3-4 days in long day conditions. Seedlings were then transferred into ½ liquid MS medium in 12-well culture plates (Thermo Fisher Scientific, Waltham, MA, US), and grown for another six or seven days. Each well contained three seedlings (pulled together as one sample), and every experiment included three independent samples from three independent wells. Seedlings were inoculated with 5×10^7^ cfu/ml of *Pto* DC3000 or *Pto* DC3000 (AvrRpt2), and samples were collected 6 hours after inoculation.

### Treatments with immune elicitors

Sterile seeds were sown on ½ solid MS medium to germinate and grown for 3-4 days in long day conditions. Seedlings were then transferred into ½ liquid MS medium in 12-well culture plates (Thermo Fisher Scientific, Waltham, MA, US), and grown for another six or seven days. Each well contained 3 seedlings (pulled together as one sample), and every experiment included 3 independent samples from 3 independent wells. The flg22 peptide or salicylic acid were added into the liquid medium to a final concentration of 100 nM and 0.5 mM, respectively. All the plates were incubated on a shaker for 5-10 minutes following addition of the elicitor. Samples were harvested into 1.5 ml tubes at different time points after treatment, as indicated in the figures.

### Root growth assay

Sterile seeds were sown on ½ solid MS medium and stratified at 4°C for 3 days. Then the plates were placed into a growth chamber for 1 day. The germinated seedlings were transferred to the new ½ solid MS medium (square plates). Square plates were placed vertically into a growth chamber and root length was measured 5 days later.

### RNA isolation, RT-PCR, and RT-qPCR

For RNA extraction, plant tissues were collected in 1.5 ml microfuge tubes with one metal bead and the tubes were immediately placed into liquid nitrogen. Samples were ground thoroughly with a TissueLyser (QIAGEN, Duesseldorf, Nordrhein-Westfalen, Germany) for 1 minute, and placed back in liquid nitrogen. Total RNA was extracted with the E.Z.N.A. Plant RNA kit (Omega Bio-Tek, Norcross, GA, US) with in-column DNA digestion and an additional sample treatment with DNAase (Omega Bio-Tek, Norcross, GA, US). First-strand cDNA was synthesized using the iScript cDNA synthesis kit (Bio-Rad, Hercules, CA, US) with a volume of 20 μl. For RT-PCR, the PCR reaction was performed in 50 μl using the Q5 Hot Start High-Fidelity DNA polymerase (New England Biolabs, Ipswich, MA, US) with 35 or 38 cycles. For quantitative RT-PCR (RT-qPCR), PCR reactions were performed in 20 μl using the iTaq Universal SYBR Green supermix kit (Bio-Rad, Hercules, CA, US) in the StepOnePlus Real-Time PCR System (Applied Biosystems, Foster City, CA, US). Data were analyzed with Excel and GraphPad Prism 6.

### Confocal microscopy imaging

Cotyledons from 3-4 day-old long day-grown Arabidopsis seedlings were imaged using a Confocal Laser Scanning Microscope (CLSW) Platform: Leica TCS SP8 (Leica, Mannheim, Germany). The GFP was excited with an argon laser at 488 nm, and its emission was detected at 500-550 nm. For plasmolysis assays, cotyledons were placed into a 1 M NaCl solution on glass slides, and GFP was observed and recorded after 5-10 minutes. For FM4-64 staining, cotyledons from 4 day-old seedlings were cut and soaked into 5 ng/|jl FM4-64 solution as described previously (Beck et al., 2012) for 5 minutes; samples were then transferred to water on glass slides and covered with coverslips. For dual channel simultaneous observation, the fluorescence signal of FM4-64 was excited with an argon laser at 561 nm and its emission was observed at 580-650 nm; the GFP was excited at 488 nm and observed at 500-550 nm. For co-localization assays, 3 week-old *N. benthamiana* leaves were co-infiltrated with *Agrobacterium* GV3101 (pMP90) carrying plasmids to express *RFP-IAN9* and *GFP-IAP1.* Leaves were co-infiltrated with the GV3101 strain carrying plasmids to express *RFP-IAN9* and *GFP* as control. Three-mm leaf discs were punched from the whole leaf 24 hours post infiltration and transferred to water on glass slides. For dual channel image acquisition, GFP and RFP were excited at 488 nm and 561nm respectively; emission was collected at 500-550 nm for GFP and 580-650nm for RFP.

### MAPK activation and western blot assays

For protein extraction, Arabidopsis seedlings or leaf discs from *N. benthamiana* were ground with a Tissue Lyser (QIAGEN, Hilden, NordrheinWestfalen, Germany). Samples were then re-suspended in lysis buffer [2x loading buffer: 100 mM Tris-HCl (pH 6.8), 10% Glycerol, 2% SDS, 0.03% bromophenol blue], boiled at 95°C for 10 minutes, and centrifuged at 14,000 g for 5 minutes before loading in SDS-PAGE acrylamide gels. Western blots were performed using anti-GFP (Sigma G6795), anti-Luciferase (Sigma L0159), anti-Mouse IgG-Peroxidase (Sigma A2554), and anti-Rabbit IgG-Peroxidase (Sigma A0545).

MAPK activation assays were performed as previously described (Macho et al, 2012), with minor modifications. Briefly, 7 day-old Arabidopsis seedlings grown on solid % MS were transferred to water and then treated with 100 nM flg22 for 10 minutes after vacuuming 5 minutes (15 minutes in total). Anti-pMAPK [Phospho-p44/42 MAPK (Erk1/2) (Thr202/Tyr204) XP Rabbit mAb; Cell Signaling 4370] was dissolved in 3% gelatin (Sigma G7041) and used to hybridize the membranes. All membraneswere stained with Ponceau stain (Sangon Biotech, Shanghai, China) to verify equal loading.

### ROS burst

ROS burst assays were conducted as previously described (Sang and Macho, 2017). Briefly, 4 mm leaf discs from 5-6 week-old Arabidopsis plants grown in short day conditions were transferred to 96-well microplates (PerkinElmer, Waltham, MA, US) with 100 μl Mili-Q water per well and incubated overnight. Water was then removed and ROS burst was elicited by adding 100 μl of a solution containing 100 ng flg22, 100 μM luminol, and 20 μg/mL horseradish peroxidase. The luminescence was recorded over 40 minutes using a Thermo Scientific VARIOSKAN FLASH (Thermo Fisher Scientific, Waltham, MA, US).

### Co-IP and large-scale IP for LC-MS/MS

Leaves from 3-4 week-old *N. benthamiana* plants were co-infiltrated with *Agrobacterium* GV3101 (pMP90) carrying plasmids to express *GFP-IAP1* and *Cluc-IAN9;* leaves co-infiltrated with GV3101 carrying plasmids to express *GFP-AIP1* and *Cluc* or *GFP* and *Cluc* were used as controls. Total proteins were extracted 24 hours later, and immunoprecipitation was performed with GFP-trap beads (Chromotek, Am Klopferspitz, Planegg-Martinsried, Germany) as described previously (Sang et al, 2016). Proteins were stripped from the beads by boiling in 50 |jl SDS loading buffer for 10 mins. Immunoprecipitated proteins were separated on SDS-PAGE acrylamide gels and western blots were performed as described above. Large-scale immunoprecipitation assays for LC-MS/MS were performed as described before (Kadota et al. 2016; Sang et al. 2016), using 5 g of 10 day-old Arabidopsis seedlings before or after treatment with 100 nM flg22 or 0.5 mM SA.

### Plasmid construction

See Table S3 for the sequence of all the primes used in this study. Free GFP fragment (with stop codon) was amplified from the plasmid pGWB505 (Nakagawa et al, 2007). The purified PCR product was transferred into entry vector pDONR/ZEO (Thermo Fisher Scientific, Waltham, MA, US) by BP reaction, and then recombined into pGWB602 (Nakagawa et al., 2007; 35S promoter, no tag) using LR reaction to yield the pGWB602-*GFP* plasmid. The CDS of *IAN9, IAN9 (ΔC27aa), IAN8, IAP1* (all with stop codon) were amplified from cDNA of whole Arabidopsis seedlings using primers containing attB1attB2 sites (Table S2). The PCR fragments were ligated into pDONR/ZEO by BP reaction, and then recombined with the binary vector pGWB606 (Nakagawa et al., 2007; 35S pro, N-sGFP) to yield the plasmids pGWB606-IAN9, pGWB606-IAN9 *(ΔC27aa)*, pGWB606-IAN8, and pGWB606-IAP1 through an LR reaction. pDONR/ZEO-*IAN9* was also recombined with the binary vector pGWB555 (35S promoter, N-mRFP) to yield the plasmid pGWB555-IAN9 through an LR reaction.

For luciferase complementation assays, the original vectors (Chen et al, 2008) were modified to make them Gateway-compatible. The binary vector pCAMBIA1300-nLUC was digested with SacI and *Sal*I and pCAMBIA1300-cLUC was digested with *KpnI* and *Sal*I. The gateway cassette was then amplified from pGWB505 with primers Nluc-F/R and Cluc-F/R (Table S1) and PCR products were cloned into the digested pCAMBIA1300 vectors using ClonExpress^®^ II One Step Cloning Kit (Vazyme Biotech, Nanjing, China) to yield the new version of the binary plasmids, pGWB-*Nluc* and pGWB-*Cluc* containing the Gateway cassette. Then, pDONR/ZEO-*IAP1* (without stop codon) and pDONR/ZEO-*IAN9* were separately recombined with pGWB-*Nluc* and pGWB-*Cluc* through an LR reaction to yield the destination vectors pGWB-C3HC4-Nluc and *pGWB-Cluc-IAN9.*

For CRISPR/Cas9-mediated mutagenesis, the detailed sites were predicted using the CRISPR Design tool (http://crispr.mit.edu/). Primer sequences can be found in the Table S3. PCR products were ligated into the pCAS9 plasmid (Feng et al., 2013) using T4 DNA ligase (New England Biolabs, Ipswich, MA, US).

### Arabidopsis transformation

For the generation of Arabidopsis transgenic lines, *Agrobacterium-mediated* transformation was performed according to the method described before (Clough and Bent, 1998). The *Agrobacterium tumefaciens* strain GV3101 (pMP90) carrying the desired plasmids (pGWB602-GFP, pGWB605-IAN9, pGWB606-IAN9, pGWB605-IAN8, pGWB606-IAN9 (ΔC-27 aa)) were cultured overnight at 28°C, then spun down and re-suspended to OD_600_=1 in 50 ml 5% sucrose and 0.03% silwet-L77 solution. The fertilized siliques from 5-6 week-old Col-0 wild type plants (grown in a 16 h light/8 h dark photoperiod) were removed before flower dipping. After dipping for 1-2 minutes, the plants were covered with PE cling wrap for 16-24 hours in the dark, and then put back to normal growth conditions until seed collection. Finally, homozygous transgenic lines were obtained after resistance selection (BASTA, 15 μg/ml; hygromycin B, 25 μg/ml) and segregation ratio calculation. For the generation of the *ian9* mutant, *A. tumefaciens* GV3101 carrying the plasmid pCas9 (35S, AtU6, Hpt)-IAN9 was used to transform Col-0 wild type Arabidopsis plants. T1 plants were selected in ½ solid MS (25 μg/ml, hygromycin B), and then transferred to soil. Genomic DNA was extracted from two-week-old soil-grown T1 plants using the CTAB method (Doyle and Doyle, 1987). *IAN9* was amplified from independent DNA samples and sequenced. The plants that possessed a double peak in the sequencing results of the target sequence were chosen for collection of T1 seeds. Independent T1 seeds were sown on selective (25 μg/ml, Hygromycin B) and non-selective ½ solid MS medium and the segregation ratio was calculated. Lines with 3:1 ratio were transferred to the soil. *IAN9* and *CAS9* genes were amplified by PCR, and plants without the *CAS9* product were selected. Homozygous CRISPR/CAS9 mutant *ian9* lines without *CAS9* background were selected for further experiments.

### *Agrobacterium*-mediated transient expression in *N. benthamiana*

*A. tumefaciens* GV3101 carrying the desired plasmid (pGWB606-*IAP1*, pGWB555-*IAN9*, pGWB-*Cluc*, pGWB-*Cluc-IAN9*, pGWB-*Nluc-IAP1*) were grown on selective LB plates (10 g Tryptone, 5 g Yeast Extract, 10 g NaCl, 15 g Agar per litre) and cultivated at 28°C for 2 days. Before infiltrating 3-4 week-old *N. benthamiana* plants, *Agrobacterium* cells were resuspended in the infiltration buffer (10 mM MgCl_2_, 10 mM MES pH 5.6, acetosyringone 150 |jM) directly from plates, and diluted to an OD600 of 0.5 or 1, depending on the expression and stability of the different proteins. Samples were collected 24 hours after *Agrobacterium* infiltration.

### Luciferase complementation assay

Luciferase complementation assays were performed as previously described (Chen et al, 2008). Three-week-old *N. benthamiana* leaves were co-infiltrated with *Agrobacterium* GV3101 (pMP90) carrying the plasmids to induce the expression of *IAP1-Nluc* and *Cluc-IAN9.* One day after inoculation, the whole leaves were cut and sprayed with luciferin solution [1mM luciferin (Sigma), 0.02% Triton X-100]. The fluorescence signal was recorded using a Lumazone 1300B (Scientific Instrument, West Palm Beach, FL, US) for 10 minutes after 5 minutes in the dark.

### Fluorescence Resonance Energy Transfer - Fluorescence-lifetime imaging microscopy (FRET-FLIM)

Three-week-old *N. benthamiana* leaves were co-infiltrated with *Agrobacterium* GV3101 (pMP90) carrying the plasmids to induce the expression of *GFP-IAP1* with *RFP-IAN9* (FRET pair: donor + acceptor); leaves infiltrated with GV3101 inducing the expression of *GFP-IAP1* with *RFP* (donor + free RFP) and *GFP-IAP1* (donor alone) were used as negative control. FRET-FLIM experiments were performed on a Leica TCS SMD FLCS confocal microscope. Six-mm leaf discs of *N. benthamiana* plants transiently co-expressing donor and acceptor were visualized one day after agroinfiltration. The lifetime of the donor (**τ**) was collected and analysed as described in (Rosas-Diaz et al., 2018).

### Chemicals

The flg22 peptide (TRLSSGLKINSAKDDAAGLQIA) was purchased from Abclonal, USA. Sequencing-grade modified trypsin was purchased from Promega (Madison, WI, USA). All other chemicals were purchased from Sigma-Aldrich (St. Louis, MO, USA) unless otherwise stated.

## Acknowledgements

We thank members of the Macho and Lozano-Durán laboratories for helpful discussions; Jianming Li, Jian-Min Zhou and Jian-Kang Zhu for sharing biological materials; and Xinyu Jian for technical and administrative assistance during this work. We thank the PSC Cell Biology core facility for assistance with confocal microscopy, and the PSC Proteomics core facility for LC-MS/MS analysis. This work was supported by the Shanghai Center for Plant Stress Biology (Chinese Academy of Sciences). Research in the Macho laboratory is also supported by the National Natural Science Foundation of China (NSFC; grant 31571973) and the Chinese 1000 Talents Program. Research in the Lozano-Durán laboratory is also supported by the National Natural Science Foundation of China (NSFC; grant 31671994) and the 100 Talents Program from the Chinese Academy of Sciences. CCM is sponsored by a CAS-TWAS President’s Fellowship for International PhD Students. TR-D is the recipient of a President’s International Fellowship Initiative (PIFI) postdoctoral fellowship (No. 2016PB042) from the Chinese Academy of Sciences. The authors have no conflict of interest to declare.

## supplementary materials

**Figure S1. Alignment of IAN proteins.**

**Figure S2. Expression of *IAN9* in *Arabidopsis* seedlings.**

**Figure S3. The C-terminal region of IAN9 is important for its PM localization.**

**Figure S4. Characterization of the GK-146B08 line with an insertion in the promoter of *IAN9.***

**Figure S5. Exact mutation site in the *ian9* mutant generated by CRISPR/Cas9.**

**Figure S6. Characterization of the *ian9* mutant and *IAN9* over-expression lines.**

**Figure S7. Transcription of *IAN8* is not affected by over-expression of *IAN9.***

**Figure S8. Plant growth and early immune responses are not affected by altered expression of *IAN9.***

**Figure S9. Seedlings of the *ian9* mutant and *IAN9* over-expression do not show developmental defects.**

**Figure S10. Flg22-triggered ROS dynamics are not affected by alterations in *IAN9* expression.**

**Figure S11. *Ian9* and *iapl* mutant lines do not have altered expression of SA-dependent defence-related genes.**

**Figure S12. Analysis of the *ian9l35S::GFP-IAN9* complementation line.**

**Figure S13. Transgenic lines overexpressing GFP-IAN9 in a wild type background show a tendency to support higher bacterial loads compared to wild type or GFP-expressing lines.**

**Figure S14. Transgenic lines overexpressing GFP-IAN9 are not affected in susceptibility to *Pto* DC3000 COR-.**

**Figure S15. Characterization of IAP1.**

**Figure S16. Cluc-IAN9 associates with GFP-IAP1 but not with free GFP in *N. benthamiana* leaves.**

**Figure S17. IAN9 and IAP1 do not interact directly in *N. benthamiana* leaves.**

**Figure S18. Characterization of the *iapl* mutant alleles.**

**Figure S19. Seedlings of the *iap1* mutant lines show slightly reduced root elongation.**

**Figure S20. *IAN9* orthologs in different plant species.**

**Figure S21. *IAP1* orthologs in different plant species.**

**Table S1. Proteins associated to GFP-IAN9 identified after affinity purification followed by LC-MS/MS analysis.**

**Table S2. IAP1 interactors according to the Arabidopsis Interactome Database.**

**Table S3. List of primers used in this study**

